# Network Preservation Reveals Shared and Unique Biological Processes Associated with Chronic Alcohol Abuse in NAc and PFC

**DOI:** 10.1101/2020.05.21.108621

**Authors:** Eric Vornholt, Mohammed Mamdani, John Drake, Gowon McMichael, Zachary N. Taylor, Silviu-Alin Bacanu, Michael F. Miles, Vladimir I. Vladimirov

## Abstract

**Background:** Excessive alcohol consumption has become a growing public health concern worldwide due to the potential development of alcohol dependence (AD). Prolonged alcohol abuse leads to dysregulation of the mesocorticolimbic pathway (MCL), effectively disrupting executive functioning and the allostatic conditioning of reward response.

**Methods:** We utilized weighted gene co-expressed network analysis (WGCNA) and network preservation using a case/control study design (*n*=35) to identify unique and shared biological processes dysregulated in AD in the prefrontal cortex (PFC) and nucleus accumbens (NAc). We used correlation and regression analyses to identify mRNA/miRNA interactions and local expression quantitative trait loci (cis-eQTL) to identify genetic regulatory mechanisms for networks significantly associated with AD.

**Results:** Network analyses revealed 6 and 3 significant mRNA modules from the NAc and PFC, respectively. Network preservation revealed immune response upregulation in both regions, whereas cellular morphogenesis/localization and cilia-based cell projection processes were upregulated only in the NAc. We observed 4 significantly correlated module eigengenes (ME) between the significant mRNA and miRNA modules in PFC, and 6 significant miRNA/mRNA ME correlations in NAc, with the mir-449a/b cluster emerging as a potential regulator for cellular morphogenesis/localization dysregulation in this brain region. Finally, we identified cis-eQTLs (37 mRNA and 9 miRNA in NAc, and 17 mRNA and 16 miRNA in PFC) which potentially mediate alcohol’s effect in a brain region-specific manner.

**Conclusion:** In agreement with previous reports, we observed a generalized upregulation of immune response processes in subjects with AD, that highlights alcohol’s neurotoxic properties, while simultaneously demonstrating distinct molecular changes in subcortical brain regions as a result of chronic alcohol abuse. Such changes further support previous neuroimaging and physiological studies that emphasize the distinct roles PFC and NAc play in the development of addictive behaviors.

## Introduction

Alcohol use disorder (AUD) is a debilitating psychiatric illness with negative health, economic, and social consequences for nearly 15.1 million affected adults worldwide [1]. AUD risk is dependent upon both genetic and environmental factors, with a heritability of 0.49 [2]. The neurobiological framework for understanding how benign, recreational alcohol use leads to AUD follows various theories [3–5], with the most commonly accepted being the cyclical model of addiction [6]. This theory provides valuable insight into the functional specialization of different brain regions that underlie behavioral maladaptations associated with AUD [7]. However, the genetic architecture and molecular mechanisms contributing to alcohol-facilitated neuroadaptations remain widely unknown.

Postmortem brain studies provide the unique opportunity to interrogate neurobiological changes associated with addiction across brain regions and neural pathways [8, 9]. Among these, the mesocorticolimbic system (MCL), which connects the ventral tegmental area (VTA) to the prefrontal cortex (PFC), and nucleus accumbens (NAc), has proven especially sensitive to alcohol-associated neuroadaptations [10–12]. Recent postmortem brain studies of AUD have focused on examining gene and noncoding RNA expression as the biological intermediate between genetic variation and molecular function [13–19]. While single gene expression differences are continuously explored, network approaches, such as weighted gene co-expressed network analysis (WGCNA), allows genes with correlated expression to cluster into modules that then can be independently analyzed to identify dysregulated biological processes and molecular pathways underlying the etiology of AUD [20]. Others and we have successfully implemented this method to identify gene networks associated with AUD within the MCL and other brain regions [16, 18]. While postmortem brain expression differences alone are insufficient to infer a causal relationship between AUD and neurobiological function, the integration of genetic information via expression quantitative trait loci (eQTL) analysis can help elucidate the regulatory mechanisms for genes whose expression is associated with AUD [21]. Studying mRNA/miRNA interactions may further reveal functional relationships that mediate the differential expression of risk AUD genes based on the role miRNAs play in the destabilization and degradation of their target genes [22].

Thus, in this study, we seek to expand upon previous research by jointly analyzing two key MCL areas, the NAc and PFC, to identify unique and shared neurobiological processes associated with alcohol dependence (AD). To achieve this, we utilize differential gene expression, WGCNA, network preservation, mRNA/miRNA interactions, and cis-eQTL analysis using a case/control study design.

## Materials and Methods

### Tissue Processing and RNA Extraction

Postmortem brain tissue from 42 AD cases and 42 controls was provided by the Australian Brain Donor Programs of New South Wales Tissue Resource Centre (NSW TRC) under the support of The University of Sydney, National Health and Medical Research Council of Australia, Schizophrenia Research Institute, National Institute of Alcohol Abuse and Alcoholism, and the New South Wales Department of Health [8]. Samples were excluded based on: (1) history of infectious disease, (2) circumstances surrounding death, (3) substantial brain damage, and (4) post-mortem interval > 48 hours (*Supplemental 1*). Total RNA was isolated from flash-frozen PFC and NAc tissue using the mirVANA-PARIS kit (Life Technologies, Carlsbad, CA) following the manufacturer’s suggested protocol. RNA concentrations and integrity (RIN) were assessed via Quant-iT Broad Range RNA Assay kit (Life Technologies) and Agilent 2100 Bioanalyzer (Agilent Technologies, Inc., Santa Clara, CA) respectively. NAc samples were matched for RIN (mean=6.9, ±0.84), age, sex, ethnicity, brain pH, and PMI as part of a previous study [18] yielding a total of 18 case-control matched pairs (n=36). As PFC samples were matched to NAc samples to ensure consistencey with our comparison analyses, we had reduced flexibility in identifying PFC samples with the best RIN numbers, leading to slightly lower RINs in our PFC samples (mean=4.5, ±2.04).

### Gene Expression Microarray and Data Normalization

Gene expression was assayed using Affymetrix GeneChip Human Genome U133A 2.0 (HG-U133A 2.0) on 22,214 probe sets spanning ~ 18,400 mRNA transcripts, and the Affymetrix GeneChip miRNA 3.0 microarray interrogating the expression of 1733 mature miRNAs as previously described [23]. None of the mRNA or miRNA probes were excluded based on quality control criteria outlined in previous studies [18]. Raw probe data were GCRMA background corrected, log_2_ transformed, and quantile normalized using Partek Genomics Suite v6.23 (PGS; Partek Inc., St. Louis, MO) to obtain relative gene expression values. A principal component analysis was used to identify potential outlier samples. Only one case sample was removed from the analyses, leaving 18 controls and 17 cases (n= 35) for both brain regions. It has become widely accepted to verify a subset of microarray-generated gene expression changes via an independent platform such as qPCR. Considering our extensive use of the Affymetrix platform in the past, and similar to other groups [24], we decided not to include microarray validation in this study, as we have already ‘validated’ the same array and platform with an independent qPCR experiment previously with a concordance between microarray and qPCR validation in the past exceeding 80% [18, 25].

### Analysis of Differential Gene Expression

The relationship between AD case status and normalized gene expression was analyzed via bidirectional stepwise regression for each gene to adjust for demographic and postmortem covariates independently. Regression coefficients were calculated in RStudio (ver. 1.1.463) using the Stats package (ver. 3.5.1); for more details, see *Supplemental 2*.

### Network Analyses

WGCNA was performed using the WGCNA package in RStudio (ver. 1.66). All nominally significant genes (p≤0.05) were used to generate a signed similarity matrix via pair-wise Pearson correlations. The nominal significance was chosen to include genes with smaller effect sizes, albeit true positive signals, exclude genes with low disease variance, i.e., likely not associated with AD, and to provide a sufficient number of genes for the network analysis. WGCNA was performed as outlined previously by us and others [16, 18] and in *Supplemental 2*. Module eigengenes (MEs), serving as a single aggregate expression value for each of the coexpressed modules, were correlated to AD case-status and available demographic/biological covariates. To validate WGCNA module clustering, we further performed a bootstrap based resampling of 100 iterations with replacement. Next, using WGCNA with the clusterRepro (ver. 0.9) package in RStudio, we identified the level of module preservation between the PFC and NAc by comparing adjacency matrices and calculating the composite preservation statistic (Z_summary_). A Z_summary_ >10 indicates strong evidence for network preservation, Z_summary_ <10 >2 indicates weak evidence of network preservation and Z_summary_ <2 indicates no module preservation, as outlined previously [26].

### Gene Set Enrichment Analysis

Gene set enrichment was performed using ShinyGo (ver. 0.61) gene annotation database [27]. Gene lists from the significant AD modules from NAc and PFC were enriched using GO biological processes consisting of 15,796 gene sets from the Ensembl BioMart release 96; all p-values for significantly enriched gene sets are FDR adjusted (FDR of 5%). We further performed cell type enrichment using the “*userListEnrichment”* option within the WGCNA package in R (ver. 1.66) as previously described [18]. Statistical significance of brain-list enrichment was determined via a hypergeometric test; all p-values were adjusted at FDR of 5%.

### Hub Gene Prioritization

Hub genes were defined based on the strength of intramodular connectedness, calculated from the absolute value of the Pearson’s correlation coefficient between module eigengene and expression values. This value is denoted as module membership (MM). Hub genes were prioritized for downstream analysis based on MM of r ≥0.80 and a significant gene correlation with AD (at p≤0.05). These hub genes may serve as promising prospective targets for future research based on their potential as driving factors for the etiology of AD within biologically relevant molecular and neurobiological pathways.

### Cis-eQTL Analysis

DNA from the postmortem brain sample was processed and genotyped as part of a larger GWAS study [18]. Genotypes with excessive missingness (greater than 20%) and monomorphic for homozygous major and minor alleles were removed. We then isolated only cis-eQTLs, which are defined as SNPs 500kb from the start/stop positions for each hub gene and significant miRNA of interest. These SNPs were subsequently pruned with Plink v1.9 to exclude variants in linkage disequilibrium (R^2^ ≥0.7). For eQTL detection, the pruned genotypic dataset and the expression of selected genes were analyzed via MatrixEQTL package (ver. 2.2) in R using a linear regression model adjusting for covariates. The effect of SNP on AD case status via gene expression was assessed by including an interaction (SNP x Diagnosis) term using the “*modelLINEAR_CROSS”* argument.

### mRNA/miRNA Target Prediction

The relationship between the miRNA and mRNA MEs from each brain region was examined via Pearson’s correlations. Significant miRNA/mRNA MEs correlations (FDR of 5%) were selected for further, more detailed, series of analyses, in which individual Hub gene expression from the selected modules are correlated to miRNA expression via Pearson’s correlations using the miRLAB package in R (ver. 1.14.3).

## Results

### AD Case/Control Differentially Expressed Genes (DEG)

A bidirectional stepwise regression revealed 3,536 and 6,401 DEG in the PFC and NAc, respectively, at the nominal p ≤0.05, of which 1,279 DEG were shared between the two regions. In both regions, 603 genes were downregulated and 494 were upregulated, whereas 182 genes from the PFC and NAc were expressed in opposite directions. After adjusting p-values (FDR of 5%), we identified 1,841 DEG from the NAc and 70 from the PFC. The miRNA regression analysis identified 430 and 170 nominally significant miRNAs in the NAc and PFC, respectively. At FDR of 5%, 168 miRNAs were differentially expressed in NAc; however, no miRNA reached FDR significance in the PFC. For detailed information about regression coefficients, the regression models used for each transcript, and frequency of covariates incorporated into the analysis, see *Supplemental 3*.

### mRNA Gene Network Module Clustering

Within NAc, we identified 23 distinct modules ranging in size from 37 transcripts (NAc_*darkolivegreen*_) to 1,584 transcripts (NAc_*blue*_) (***Figure 1A***). At Bonferroni adjusted p≤0.05, we identified 6 significant modules from the NAc after correlating module MEs with AD case status. Among these, NAc_*darkgreen*_ was the only significantly downregulated module, whereas NAc_*darkorange*_, NAc_*purple*_, NAc_*magenta*_, NAc_*skyblue*_, and NAc_*greenyellow*_ were all significantly upregulated in AD cases relative to controls (***Figure 1B***). In PFC, we identified 17 co-expressed modules ranging in size from 37 transcripts (PFC_*darkturquoise*_) to 889 transcripts (PFC_*turquoise*_) (***Figure 1C***), of which three modules were significantly correlated to AD at Bonferroni adjusted p≤0.05. Of these, the PFC_*pink*_ module was significantly downregulated, while PFC_*darkred*_ and PFC_*lightgreen*_ were significantly upregulated among AD cases (***Figure 1D***). The bootstrap validation of such identified networks showed consistent module clustering when compared to the original gene networks *(Supplemental 4)*.

**Figure 1:**
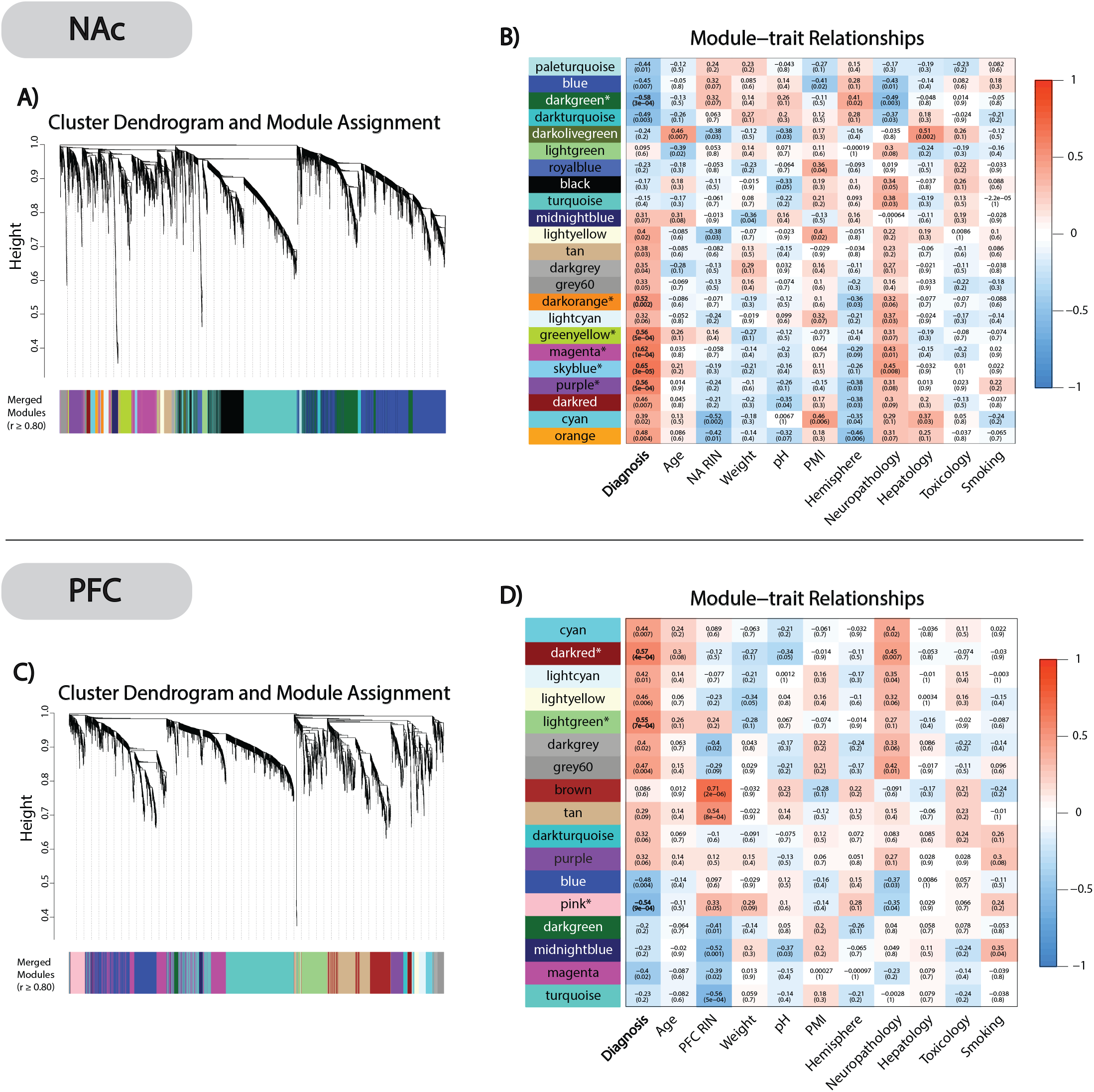
WGCNA clustering and module-trait relationships. ***A)*** NAc cluster dendrogram and module assignment with dissimilarity based on topological overlap. The 6,401 selected transcripts were clustered into 23 distinct modules. ***B)*** NAc module-trait relationship heatplot correlating (Pearson’s) module MEs with AD diagnosis and covariates. Uncorrected p-values are givin in parenthesis below each correlation coefficient. 6 AD associated significant modules (NAc_*darkgreen*_, NAc_*darkorange*_, NAc_*purple*_, NAc_*magenta*_, NAc_*skyblue*_, and NAc_*greenyellow*_) were identified after Bonferroni correcting p-values (*= p≤0.05). ***C)*** PFC cluster dendrogram and Module assignment. The 3,536 selected transcripts were clustered into 17 different co-expressed modules. ***D)*** PFC module-trait relationship heatplot created as previously described. We identified 3 AD associated modules (PFC_*pink*_, PFC_*darkred*_, and PFC_*darkgreen*_) after Bonferroni correcting p-values (*= p≤0.05).

### NAc and PFC Network Preservation

In our next step, we performed a network preservation analysis to determine how well co-expressed networks from the PFC are conserved in NAc and vice versa. We focused primarily on the *Z*_*summary*_ and *Median Rank* network preservation statistics because Z_*summary*_ estimates network overlap by also taking into consideration network connectivity. *Median Rank* being invariant to module size, provides a more accurate estimate of network preservation, i.e., larger networks tend to be more conserved due to their size alone. Based on these analyses, we observed that NAc_*darkorange*_ and NAc_*purple*_ show little to no network preservation (Z_summary_ <2), NAc_*skyblue*_, NAc_*darkgreen*_, PFC_*darkred*_, and PFC_*pink*_ show moderate levels of network preservation (2< Z_summary_ <10), and NAc_*greenyellow*_, NAc_*magenta*_, and PFC_*lightgreen*_ showed high levels of network preservation (Z_summary_ >10) (***fig. 2A/B***). For detailed information about the individual density and connectivity statistics that were used to create the composite network preservation statistics, see the *Supplemental 5*.

**Figure 2:**
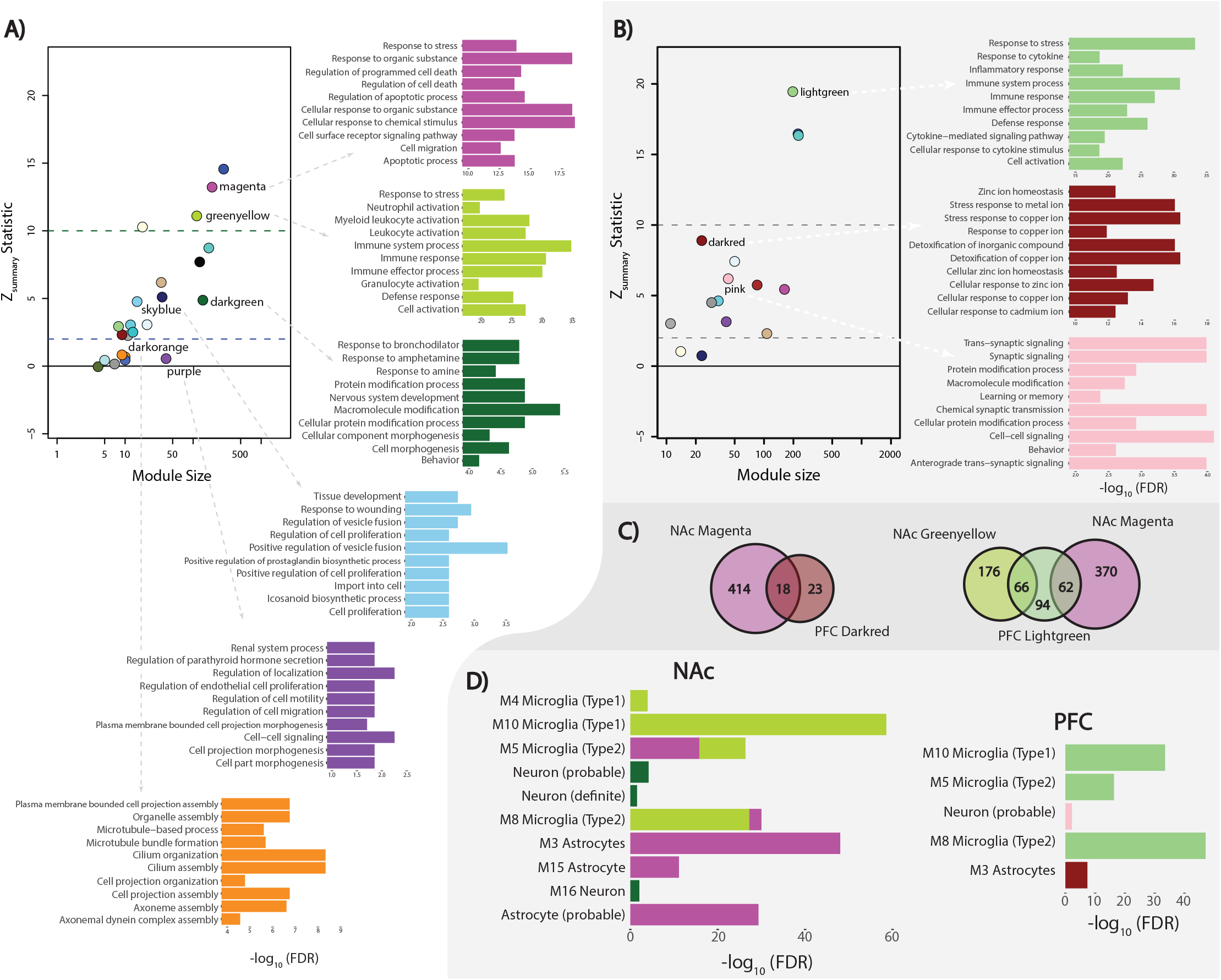
Network preservation and gene-set enrichment. ***A)*** NAc Z-summary statistic calculated as an aggregate of network preservation statistics (*Preservation level:* high = *Z*>10; moderate *=* 2<*Z*<10; low = Z<2) with color corresponded top-10 most significant (−log_10_(FDR) transformed) GO biological processes for significant AD associated modules. ***B)*** PFC Z-summary statistic and corresponding GO biological processes term (−log_10_(FDR) transformed). ***C)*** Venn-diagram of the shared transcripts from highly preserved NAc modules (NAc_*magenta*_ and NAc_*greenyellow*_) and their corresponding significant PFC modules (PFC_*lightgreen*_ and PFC_*darkred*_). ***D)*** Brain cell type gene-set enrichment from the NAc and PFC (−log_10_(FDR) transformed). Colors correspond with their respective modules (NAc_*greenyellow*_, NAc_*magenta*_, NAc_*darkgreen*_, PFC_*pink*_, PFC_*darkred*_, and PFC_*lightgreen*_) with single gene sets enriched in modules.

### Identifying Biological Processes and Cell-type Enrichment of the AD Significant Modules

To gain perspective on the biological underpinnings of the significant gene networks from NAc and PFC, we performed a gene-set enrichment analysis, GO biological processes annotation (ShineyGO ver.61) and neuronal cell type enrichment for the two regions. Given the core aim of this study to identify unique and shared gene networks associated with AD in NAc and PFC, we focused on NAc modules highly (i.e., NAc_*greenyellow*_ and NAc_*magenta*_) and poorly (i.e., NAc_*darkorange*_, and NAc_*purple*_) preserved in PFC, respectively. The preserved NAc modules (NAc_*greenyellow*_ and NAc_*magenta*_) are primarily associated with the immune response process (FDR ≤0.05) believed to be a consequence of neurotoxicity caused by chronic alcohol abuse (***Figure 2A***). Additionally, these modules are enriched among microglia and astrocyte cell types (FDR ≤0.05), which is expected based on the functional properties of these glial cells (***Figure 2D***). The poorly preserved NAc modules are enriched within gene-sets associated with cilia-based cell projection and cell morphogenesis (FDR≤0.05) (***Figure 2A***).

In an attempt to identify highly preserved PFC modules within NAc, we performed additional gene-set enrichment analysis on the PFC modules associated with AD (***Figure 2B***). Similar to NAc_*greenyellow*_ and NAc_*magenta*_ modules, we see the highly preserved PFC_*lightgreen*_ module to be associated with immune response processes (FDR ≤0.05) and significant microglial cell type enrichment (FDR ≤0.05) (***Figure 2D***). The relatively small PFC_*darkred*_ module and NAc_*magenta*_, were moderately preserved with each other (***Figure 2C***) with PFC_*darkred*_ also showing astrocyte cell type enrichment (***Figure 2D***). Interestingly, the genes in one family of immune response proteins, metallothioneins (MTs), were differentially expressed in both brain regions between cases and controls (***Figure 3***). For the complete gene set enrichment analyses for all modules associated with AD in NAc and PFC see *Supplemental 6*.

**Figure 3:**
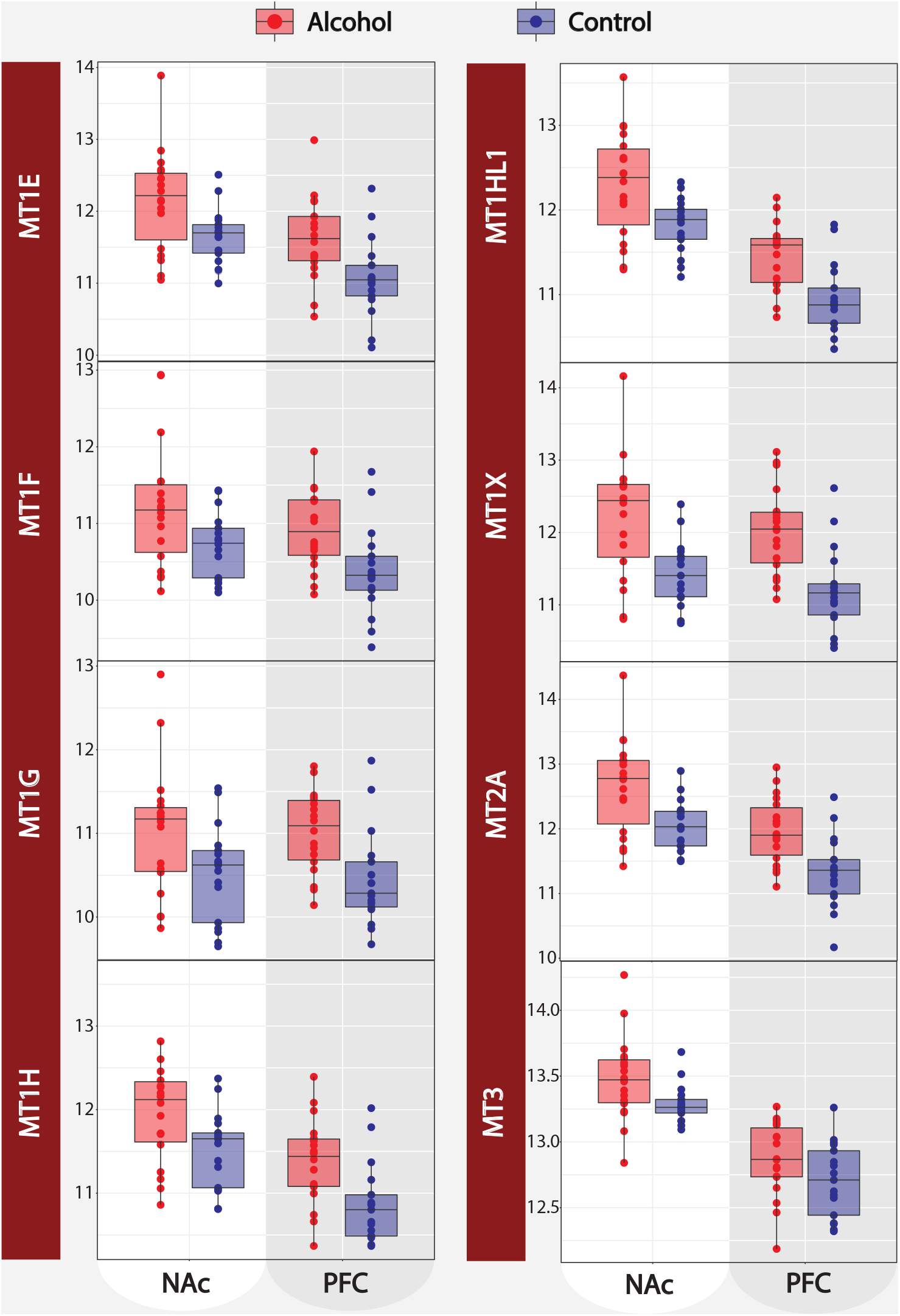
Metallothionine gene expression. Relative expression of 8 metallothionein cluster genes (*MT1E, MT1F, MT1G, MT1H, MT1HL1, MT1X, MT2A*, and *MT3*) comparing AD case/control status for both the NAc and PFC.

### Hub Genes of Potential Biological Significance

To identify candidate hub genes of potential biological significance, we focused on the relationship between intramodular connectivity (i.e., module membership (MM)) and gene significance (GS) to AD case status. Of the 459 genes from the 3 significant PFC modules (PFC_*darkred*_, PFC_*lightyellow*_, and PFC_*pink*_), we identified a total of 99 unique hub genes with MM ≥0.80. Our analysis of the 1843 genes in the 6 significant modules in NAc revealed a total of 433 unique candidate hub genes (MM ≥0.80). For full PFC and NAc transcript information regarding individual MM and GS see *Supplemental 7*.

### Detection of miRNA Gene Network Modules in NAc and PFC

In NAc and PFC, we identified co-expressed miRNA modules with varying levels of significant association to AD case status. The NAc miRNA data revealed 430 nominally significant loci, which clustered in 5 co-expressed modules ranging from 18 (NAcmi_*green*_) to 259 (NAcmi_*turquoise*_) loci in size, of which, we identify three significantly correlated with AD modules (NAcmi_*yellow*_, NAcmi_brown_, and NAcmi_*turquiose*_) at Bonferroni adjusted p≤0.05. Of these NAcmi_*yellow*_ and NAcmi_*brown*_ were downregulated, whereas NAcmi_*turquoise*_ upregulated within AD cases relative to controls (***Figure 4A***). The 170 miRNA transcripts from the PFC clustered into 6 modules ranging in size from 9 (PFCmi_*red*_) to 55 miRNA transcripts (PFCmi_*turquoise*_), of which PFCmi_*yellow*_ and PFCmi_*red*_, remain significant at Bonferroni adjusted p≤0.05; both miRNA modules were downregulated in the AD cases (***Figure 4D***). For more detailed information on the miRNA WGCNA from both brain regions, including individual transcript MM and GS values, please refer to *Supplemental 8*.

**Figure 4:**
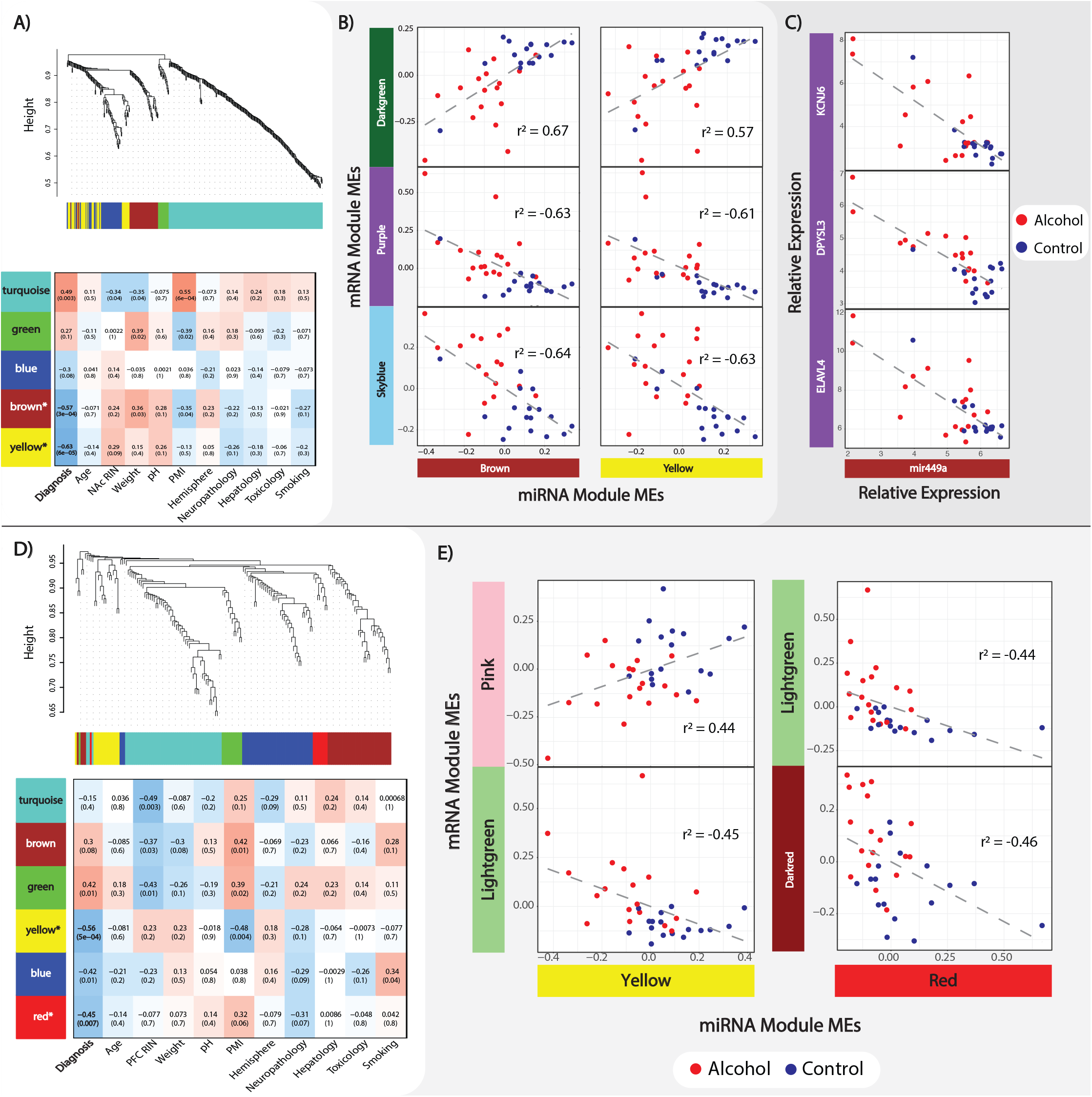
MiRNA WGCNA and mRNA:miRNA interaction. ***A)*** Nac miRNA cluster dendrogram and module assignment with module trait relationship heat map, both as previously described in *Figure 1*. ***B)*** Bonferroni adjusted significant (p≤0.05) NAc mRNA/miRNA module ME correlations (Pearson’s). Alcohol and control groups are separated by color to emphasize correlation clustering. ***C)*** Significant (FDR≤.05) correlation (Pearson’s) between mir-449a and selected mRNA transcripts from the low network preserved NAc_*purple*_ module. ***D)*** PFC miRNA cluser dendogram Module assignment along with module-trait relationship heatmap. ***E)*** Bonferroni adjusted significant (p≤0.05) NAc mRNA:miRNA module ME correlations (Pearson’s).

### MiRNA Networks Show Unique Patterns of Regulation

Within the NAc, we identified 2 significant positive mRNA/miRNA ME correlations and 4 negative ME correlations at Bonferroni adjusted p≤0.05 (***Figure 4B***). When individual hub genes from the significant AD mRNA modules were correlated to the significant AD miRNA modules, we identified 1,801 significant mRNA/miRNA interactions (FDR of 1%) spanning 318 genes and 68 miRNA loci (*Supplemental 9*). Interestingly, we observed 97% (35/36) of the purple mRNA module hub genes to be negatively correlated with either mir-449a or mir-449b from NAcmi_*brown*_. (FDR of 1%) (***Figure 4C***). In PFC, we identified one positive mRNA/miRNA ME correlation and 3 negative correlations at Bonferroni adjusted p≤0.05 (***Figure 4E***). Individual mRNA/miRNA interaction analysis from the PFC revealed 6 mRNA/miRNA interactions (FDR of 1%) spanning 6 genes and one miRNA transcript, mir-485-5p. For a full list of individual mRNA/miRNA interactions, see *Supplemental 9*.

### Brain Region Specific cis-eQTL Regulation of Differential Gene Expression

In NAc, we detected a total of 36 mRNA eQTLs spanning 17 unique genes and 9 miRNA eQTLs covering 4 different miRNA (FDR ≤0.10). Of the 17 hubs with significant eQTLs, 7 are from NAc_*darkgreen*_ (*VRK1, INPP4A, HMP19, DKK3, PCDH8, RNF34,* and *RASGRP1*), 4 from NAc_*greenyellow*_ (*FCGR3A, CTSS, AASS,* and *RNASE4*), 3 from NAc_*darkorange*_ (*DNALI1, CCDC81*, and *SPAG6*), 2 from NAc_*purple*_ (*HIVEP1* and *GNAS*), and one from NAc_*magenta*_ (*VAMP5*). Within the PFC we identified 34 eQTLs spanning 16 unique genes and 18 miRNA covering 7 different miRNA transcripts (FDR ≤0.10). Of these, 11 genes are from PFC_*lightgreen*_ (*SERPINH1, CDKN1A, PNP, EMP1, FKBP5, IL4R, TNFRSF10B, RTEL1/TNFRSF6B, SERPINA1, MAFF,* and *SERPINA2*) and 5 from PFC_*pink*_ (*GAD2, ACTL6B, KCNF1, SEZ6L,* and *EFNB3*). We selected specific hub gene eQTLs from the highly conserved NAc_*greenyellow*_ module and poorly conserved NAc_*darkorange*_ to demonstrate that the genetic impact on gene expression can be associated with clinical status and specific brain regions (***Figure 5***). For the full list of cis-eQTL, please refer to *Supplemental 10*.

**Figure 5:**
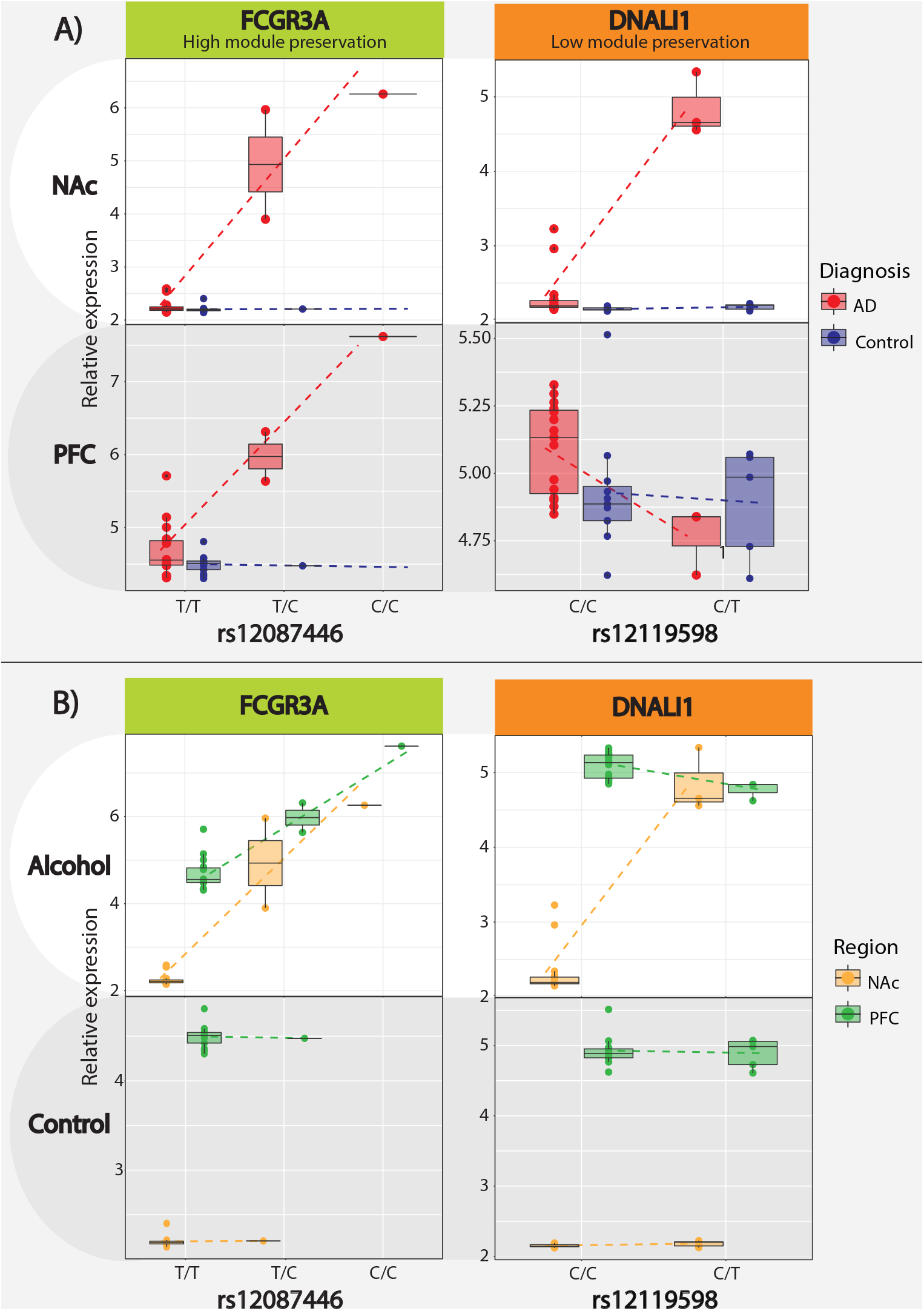
Cis-eQTL Analysis. ***A)*** Cis-eQTL box plot directly comparing AD case/control designation with the *FCGR3A:rs12087446* eQTL from the high network preservation NAc_*greenyellow*_/PFC_*lightgreen*_ module and the *DNALI1:rs12119598* eQTL from the low preservation NAC_*darkorange*_ module. ***B)*** Alternative boxplot visualization of the same cis-eQTL directly comparing differences between brain regions.

## Discussion

AUD continues to be a growing public health concern with a complex and poorly understood etiology as recreational alcohol use becomes habitual and problematic. The broad goal of this study is to identify potential neurobiological processes associated with chronic alcohol use via analyzing brain region-specific gene networks from the NAc and PFC. To understand the uniquely human nuances of addiction, it is important to investigate how chronic alcohol use impacts expression changes in the evolutionarily newer cortical areas, in contrast to the older, more evolutionarily conserved subcortical brain regions [28]. Here, we attempt to achieve that by teasing apart the neurobiological dichotomy between alcohol specific reward conditioning in the NAc and executive function disruption in the PFC [6]. In contrast to previous reports from our group [18], in this study, we tailored our analyses mainly to compare region-specific gene networks with high and low levels of conservation between NAc and PFC.

Our network analyses are consistent with previous reports from our lab and others, showing the upregulation of immune response mechanisms among AD cases as a byproduct of alcohol’s neurotoxic effects [29]. The immune-related modules show significant enrichment for both astrocyte and microglial cell types, which is not only consistent with previous alcohol studies but is also validated by known astrocyte and microglia immune function in the brain [30, 31]. More importantly, we observed generalized up-regulation of immune response mechanisms within both the PFC and NAc, suggesting that the neurotoxic response to chronic alcohol use is ubiquitous across cortical and subcortical brain regions. Interestingly, in both PFC and NAc, we further identified differentially expressed genes in the metallothionine cluster (*MT1HL1*, *MT1H*, *MT1X*, *MT1E*, *MT1G*, *MT1F*, *MT2A*, and *MT3*). The metallothionine cluster is primarily responsible for maintaining the cellular homeostasis of zinc and copper ions while also regulating oxidative stress [32]. Zinc is an essential catalytic cofactor for alcohol metabolism via alcohol dehydrogenase [33]. Free or “chelated” zinc ions (Zn^2+^) are seen in abundance in the brain, specifically at ionotropic glutamate receptors such as the NMDA receptor family. The interaction between Zn^2+^ and NMDAR activity has shown to be an important contributor to synaptic plasticity through regulating postsynaptic density assembly [34]. It is well understood that chronic alcohol abuse leads to varying degrees of organ-wide zinc deficiency [35], but the neurobiological consequences of how zinc deficiency in the brain contributes to AD etiology is poorly understood. The upregulation of brain metallothionines in response to the alcohol oxidative damage may contribute to this zinc homeostatic imbalance, altering synaptic signaling cascades and effectively the behavior. We believe this interaction between chronic alcohol abuse, metallothionine expression, zinc deficiency, and synaptic plasticity is an important avenue for future research that should be explored.

In addition to identifying dysregulated immune response mechanisms, we validate findings of differential expression among signaling and neurodevelopmental processes within AD cases [13, 15, 16, 18]. However, these processes are less conserved between cortical and subcortical regions, likely due to the different neuronal composition and functional properties of the PFC and NAc [36]. Interestingly, two NAc modules that primarily associate with cilium assembly (NAc_*darkorange*_) and cellular localization/morphogenesis (NAc_*purple*_) show limited network preservation within the PFC. While CNS primary cilia have been studied in the context of special sensory cell types (i.e., associated with olfaction, gustation, hearing, and vision) [37], there is increasing evidence to suggest primary cilia is also important for adult neurogenesis [38]. Of the cilium assembly enriched genes from NAc_*darkorange*_, three of them are associated with axonemal dynein assembly (*DNAAF1*, *DNAI2*, and *DNALI1*). A recent gene expression profiling study on adolescent rat hippocampus identified increased expression of two dynein associated genes (*dnai1* and *dnah5*) [39]. Our results with recent findings suggest that cilia, and axonemal dynein assembly, may be important for understanding the neurobiological impact of alcohol use specifically in the NAc and other subcortical pathways.

Other interesting findings arise from our mRNA/miRNA interactions, e.g., when correlating MEs from mRNA and miRNA modules, we see distinct patterns between cases and controls within both brain regions. Based on the known function of miRNAs in regulating the expression of target mRNAs [22] we can infer these significant miRNA networks may serve as a driving contributor for differential network expression between AD cases and controls. Specifically, 97% (35/36) of the hub genes from the NAc_*purple*_ module are significantly negatively correlated with either mir-449a or mir-449b. Mir-449a/b have primarily been studied in the context of spermatogenesis and cellular proliferation in cancer [40–42]. Based on the mRNA-miRNA correlations, our study suggests that mir-449a/b cluster has additional functions related to cellular proliferation in the brain. Among the genes correlated with mir-449a in the NAc, *ELAVL4*, *DPYSL3*, and *KCNJ6* have shown significant associations with AD in other gene expression and genetic association studies [19, 43, 44].

In an attempt to understand the causal nature of the gene networks associated with AD, we integrated genetic information via eQTL analysis. We were able to detect a significant number of mRNA and miRNA cis-eQTLs from both brain regions. We selected highly significant eQTLs (*FCGR3A* (Fc fragment of IgG receptor IIIa)*:rs12087446* and *DNALI1* (dynein axonemal light intermediate chain 1)*:rs12119598*) to highlight the interaction between AD case status and eQTL while also demonstrating brain region-specific eQTL variation. *FCGR3A* is one of the low-affinity Fc receptor genes important for NK cell-mediated antibody-dependent cytotoxicity [45] and a hub gene from our highly conserved NAc_*greenyellow*_ and PFC_*lightgreen*_ modules. The consistent patterning of the *FCGR3A:rs12087446* eQTL between both brain regions suggests the genetic impact on chronic alcohol abuse associated immune response dysregulation might also be ubiquitous across the brain. Differential FCGR3A expression was recently shown to be associated with both alcohol preference and binge-like behaviors within the ventral tegmental area of rats [46]. In contrast, the previously mentioned *DNALI1,* a hub gene from the cilium assembly enriched NAc_*darkorange*_ module, is under the genetic control of specific eQTL only in NAc but not in PFC, highlighting the possibility that alcohol related changes to cilia organization processes are under brain region specific genetic control.

## Conclusion

The strength of this study lies in our ability to compare and contrast expression changes between subjects with AD and controls within two different brain regions. We successfully identified gene networks and biological processes from both brain regions that were validated by previous AD studies as well implicated a novel biological process (cilia assembly) and gene family (metallothionine cluster) as potentially important for the development of AD. Our mRNA/miRNA interaction analysis pinpointed mir-449a/b cluster as an important regulator of differentially expressed genes between AD cases and controls. Finally, we provided evidence that mRNA and miRNA expression differences between AD cases and controls may be regulated by brain region specific eQTLs. While our small sample size could be perceived as a limitation, by applying network approaches, we aim to mitigate this by aggregating differentially expressed genes into biologically relevant modules that can then be analyzed and interpreted within a multivariate framework, thus, effectively increasing our power to detect significant associations with AD. As part of a previous study [18], the NAc samples were matched for known confounding postmortem covariates (i.e. RIN) to decrease confounding. In the future, by gathering larger postmortem samples, we hope to increase the power as well as extend our research strategy to include additional AD etiologically relevant brain regions to validate and expand gene discovery potential. While this study provides a large set of differentially expressed genes and gene networks associated with AD, functional multi-omic studies using single cell/nuclei technology and human tissue manipulation are necessary to elucidate the complex causal nature of genetic predisposition, chronic alcohol use, gene expression and MCL associated behavioral maladaptation.

## Acknowledgments and Disclosures

This work was supported by the National Institute on Alcohol Abuse and Alcoholism (R21AA022749 awarded to Dr. Vladimirov and P50AA022537 awarded to Dr. Miles).

The authors report no biomedical financial interests or potential conflicts of interest.

## SUPPLEMENTAL LEGENDS

**Supplemental 1:** Sample demographics.

**Supplemental 2** : Detailed overview of bidirectional stepwise regression and WGCNA methodology.

**Supplemental 3:** Stepwise regression coefficients, models and covariate frequency counts.

3.1) mRNA

3.2) miRNA

**Supplemental 4:** 100 permutation robust WGCNA dendrogram clustering.

**Supplemental 5:** Network preservation Z-summary table and supplemental network preservation statistics.

**Supplemental 6:** Full GO biological processes annotation for each AD associated module from the NAc and PFC.

6.1) NAc

6.2) PFC

**Supplemental 7:** mRNA WGCNA module membership (MM) and gene significance(GS) values with AD associated modules isolated in separate tabs.

7.1) NAc

7.2) PFC

**Supplemental 8:** miRNA WGCNA module membership (MM) and gene significance(GS) values with AD associated modules isolated in separate tabs.

8.1) NAc

8.2) PFC

**Supplemental 9:** Top mRNA/miRNA correlations (NAc = top 2000; PFC = Top 500)

**Supplemental 10:** cis-eQTL analysis results.

