## Supplemental 2 for "Network Preservation Reveals Shared and Unique Biological Processes Associated with Chronic Alcohol Abuse in NAc and PFC"

Supplemental methods

**Bi-directional Stepwise Regression:**

To analyze differential gene expression between our AD case/control groups we utilized RStudio (ver. 1.1.463) with the Stats package (ver. 3.5.1) to perform a bi-directional stepwise regression for both mRNA and miRNA normalized expression data for both the NAc and PFC. The bidirectional stepwise regression analysis cycles through all available covariates (i.e. age, RIN, pH, PMI, brain weight, hemisphere, toxicology, hepatology, neuropathology status, and smoking) to identify the best-fitting model with the lowest Akaike information criteria (AIC) for each transcript. Along with regression coefficients, we identified the number of times a covariate was incorporated into a regression model and the best-fitting model for each transcript.

**WGCNA:**

Next, the similarity matrix was raised to a power (mRNA β = 14; miRNA β = 6) to approximate the scale-free topography of the adjacency matrix, in which stronger correlations are emphasized over weaker ones. Transcript interconnectedness was determined from the calculated topological overlay measure (TOM). For module detection, unsupervised hierarchical clustering (default) was used to partition modules at specified dendrogram branch cut sites using the Dynamic Tree Cut method. Highly correlated modules were then merged based on minimum merge height of r^2^ = .8 and minimum module size of 35. Conventional colors were used to categorically label co-expressed networks and the sum of relative expression within each module is represented as a single value (module eigengene) for downstream phenotypic analysis.
