## Supplemental 4 for "Network Preservation Reveals Shared and Unique Biological Processes Associated with Chronic Alcohol Abuse in NAc and PFC"

**rWGCNA:**

To ensure network robustness and minimize the potential effect of outlier samples on network structure, we used the robust ‘bootstrapped’ version of WGCNA (rWGNCA). We performed 100 iterations in which networks were created after randomly subsetting 2/3 of the total sample, as previously suggested [26]. The resulting 100 networks were merged into one large, final consensus network with the individual sub-networks showing reasonably high consistency with the final networks.

NAc

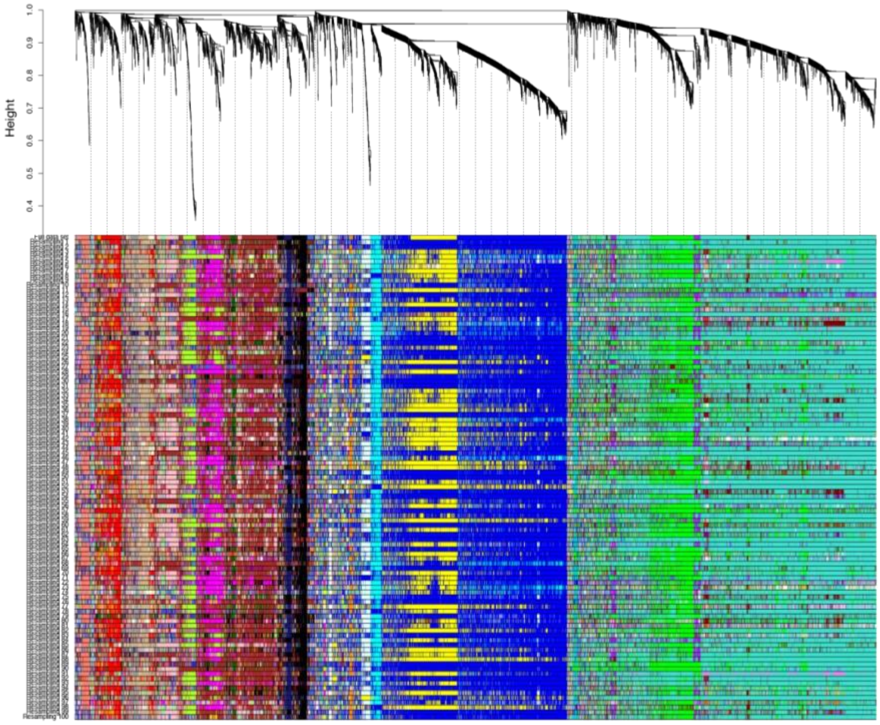


PFC

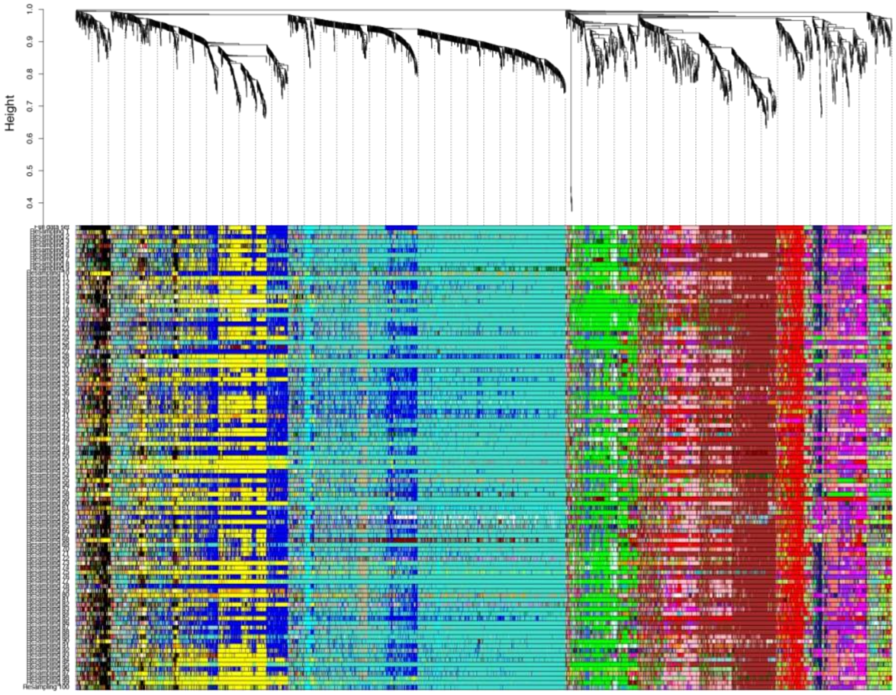
